# Estimating the effect of the 2005 change in BCG policy in England: A retrospective cohort study

**DOI:** 10.1101/567511

**Authors:** Sam Abbott, Hannah Christensen, Nicky Welton, Ellen Brooks-Pollock

## Abstract

**Background:** In 2005, England changed from universal Bacillus Calmette–Guérin (BCG) vaccination of school-age children to targeted BCG vaccination of high-risk children at birth.

**Methods:** We combined notification data from the Enhanced Tuberculosis Surveillance system, with demographic data from the Labour Force Survey to construct retrospective cohorts of individuals in England relevant to both the universal, and targeted vaccination programmes between Jan 1, 2000 and Dec 31, 2010. For each cohort, we estimated incidence rates over a 5 year follow-up period and used Poisson and Negative Binomial regression models in order to estimate the impact of the change in policy on TB.

**Results:** In the non-UK born, we found evidence for an association between a reduction in incidence rates and the change in BCG policy (school-age IRR: 0.74 (95%CI 0.61, 0.88), neonatal IRR: 0.62 (95%CI 0.44, 0.88)). We found some evidence that the change in BCG policy was associated with a increase in incidence rates in the UK born school-age population (IRR: 1.08 (95%CI 0.97, 1.19)) and weaker evidence of an association with a reduction in incidence rates in UK born neonates (IRR: 0.96 (95%CI 0.82, 1.14)). Overall, we found that the change in BCG policy was associated with directly preventing 385 (95% CI −105, 881) TB cases.

**Conclusions:** Withdrawing universal vaccination at school-age and targeting BCG vaccination towards high-risk neonates was associated with reduced incidence of TB in England. This was largely driven by reductions in the non-UK born. There was a slight increase in UK born school-age cases.

**Key Messages:**

- There is little existing literature on the impact of withdrawing universal school-age BCG vaccination and introducing high-risk neonatal BCG vaccination on TB incidence rates in the populations directly affected by the vaccination programmes.
- There was strong evidence that the change in policy was associated with a decrease in TB incidence rates in non-UK born neonates and school-age children. In the UK born individuals, there was some evidence that the change in policy was associated with an increase in TB incidence rates in those relevant to the universal school-age scheme, with little evidence of a decrease in incidence rates in those relevant to the high-risk neonatal vaccination scheme.
- Overall the change in vaccination policy was associated with preventing TB cases, mainly in the non-UK born.
- These results provide an important evaluation of the direct effects of both withdrawing and implementing a BCG vaccination programme in a low incidence, high income, country and are relevant to several other countries that have made similar changes to their vaccination programmes.

## INTRODUCTION

In 2005 England changed its Bacillus Calmette-Guérin (BCG) vaccination policy against tuberculosis (TB) from a universal programme aimed at 13 and 14 year olds to a targeted programme aimed at high-risk neonates. High risk babies are identified by local TB incidence and by the parents’ and grandparents’ country of origin. The change in policy was motivated by evidence of reduced TB transmission,(1–3) and high effectiveness of the BCG vaccine in children,(4–6) and variable effectiveness in adults.(7) Little work has been done to evaluate the impact of this change in vaccination policy.

Globally, several countries with low TB incidence have moved from universal vaccination, either of those at school-age or neonates, to targeted vaccination of neonates considered at high-risk of TB.(8) In Sweden, which discontinued universal vaccination of neonates in favour of targeted vaccination of those at high risk, incidence rates in Swedish-born children increased slightly after the change in policy. (9) In France, which also switched from universal vaccination of neonates to targeted vaccination of those at high-risk, a study found that targeted vaccination of neonates may have reduced coverage in those most at risk.(10)

The number of TB notifications in England increased from 6929 in 2004 to 8280 in 2011 but has since declined to 5137 in 2017.(1) A recent study found that this reduction may be linked to improved TB interventions.(11) Directly linking trends in TB incidence to transmission is complex because after an initial infection an individual may either develop active disease, or enter a latent stage which then may later develop into active disease. Incidence in children is a proxy of TB transmission, because any active TB disease in this population is attributable to recent transmission. Using this approach it is thought that TB transmission has been falling in England for the last 5 years, a notion supported by strain typing.(1) However, this does not take into account the change in BCG policy, which is likely to have reduced incidence rates in children.

Although the long term effects of BCG vaccination such as reducing the reactivation of latent cases and decreasing on-wards transmission are not readily detectable over short time scales the direct effects of vaccination on incidence rates can be estimated in vaccinated populations, when compared to comparable unvaccinated populations.(12) Here, We aimed to estimate the impact of the 2005 change in BCG policy on incidence rates, in both the UK and non-UK born populations, directly affected by it.

## METHODS

### Data sources

Data on all notifications from the Enhanced Tuberculosis Surveillance (ETS) system from Jan 1, 2000 to Dec 31, 2015 were obtained from Public Health England (PHE). The ETS is maintained by PHE, and contains demographic, clinical, and microbiological data on all notified cases in England. A descriptive analysis of TB epidemiology in England is published each year, which fully details data collection and cleaning.(1)

We obtained yearly population estimates from the April to June Labour Force Survey (LFS) for 2000-2015. The LFS is a study of the employment circumstances of the UK population, and provides the official measures of employment and unemployment in the UK. Reporting practices have changed with time so the appropriate variables for age, country of origin, country of birth, and survey weight were extracted from each yearly extract, standardised, and combined into a single data-set.

### Constructing Retrospective cohorts

We constructed retrospective cohorts of TB cases and individuals using the ETS and the LFS. TB cases were extracted from the ETS based on date of birth and date of TB notification.

Cohort 1: individuals aged 14 between 2000 and 2004, who were notified with TB whilst aged between 14 and 19 years old.

Comparison cohort 1: individuals aged 14 between 2005 and 2010, who were notified with TB whilst aged between 14 and 19 years old.

Cohort 2: individuals born between 2005 and 2010, who were notified with TB whilst aged 0-5 years old

Comparison cohort 2: individuals born between 2000 and 2004, who were notified with TB whilst aged 0-5 years old

Each cohort was stratified by UK birth status, with both non-UK born and UK born cases assumed to have been exposed to England’s vaccination policy. Corresponding population cohorts were calculated using the LFS population estimates, resulting in 8 population level cohorts, each with 5 years of follow up (Table 1).

**Table 1:** Summary of relevance and eligibility criteria for each cohort.

| Cohort | Vaccination programme | Eligible for the programme* | Birth status | Age at study entry | Year of study entry |
| --- | --- | --- | --- | --- | --- |
| Cohort 1 | Universal | Yes | UK born | 14 | 2000-2004 |
| Comparison cohort 1 | Universal | No | UK born | 14 | 2005-2010 |
| Cohort 1 | Universal | Yes | Non-UK born | 14 | 2000-2004 |
| Comparison cohort 1 | Universal | No | Non-UK born | 14 | 2005-2010 |
| Comparison cohort 2 | Targeted | No | UK born | Birth | 2000-2004 |
| Cohort 2 | Targeted | Yes | UK born | Birth | 2005-2010 |
| Comparison cohort 2 | Targeted | No | Non-UK born | Birth | 2000-2004 |
| Cohort 2 | Targeted | Yes | Non-UK born | Birth | 2005-2010 |

\* Eligible signifies that the cohort fit the criteria for the programme and entered the study during the time period it was in operation not that the cohort was vaccinated by the programme.

### Statistical methods

We estimated incidence rates (with 95% confidence intervals) by year, age and place of birth as (number of cases) divided by (number of individuals of corresponding age). UK birth status was incomplete, with some evidence of a missing not at random mechanism. We imputed the missing data using a gradient boosting method (see supplementary information). We then used descriptive analysis to describe the observed trends in age-specific incidence rates over the study period, comparing incidence rates in the study populations relevant to both vaccination programmes before and after the change in BCG policy.

We calculated Incidence Rate Ratios (IRRs) for the change in incidence rates associated with the change in BCG vaccination policy (modelled as a binary breakpoint at the start of 2005) for both the UK born and non-UK born populations that were relevant to the universal programme, and for the targeted programme using a range of models. We considered the following covariates: age,(1, 7) incidence rates in both the UK born and non-UK born who were not in the age group of interest,(1) and year of study entry (as a random intercept). We first investigated a univariable Poisson model, followed by combinations of covariates (supplementary table S1). We also investigated a Negative Binomial model adjusting for the same covariates as in the best fitting Poisson model. The models were estimated with a Bayesian approach using Markov Chain Monte Carlo (MCMC), with default weakly informative priors (see supplementary information). Model fit, penalised by model complexity, was assessed using the leave one out cross validation information criterion (LOOIC) and its standard error.(13) Models were ranked by goodness of fit, using their LOOIC, with a smaller LOOIC indicating a better fit to the data after adjusting for the complexity of the model. No formal threshold for a change in the LOOIC was used, with changes in the LOOIC being evaluated in the context of their standard error. The inclusion of the change in policy in the best fitting model was tested by refitting the model excluding the change in policy and estimating the improvment in the LOOIC. Once the best fitting model had been identified we estimated the number of cases prevented, from 2005 until 2015, for each vaccination programme in the study population relevant to that programme (see supplementary information).

### Implementation

R 3.5.0 was used for all analysis.(14) Missing data imputation using a GBM was implemented using the h2o package.(15) Incidence rates, with 95% confidence intervals, were calculated using the epiR package.(16) The brms package,(17) and STAN,(18) was used to perform MCMC. Models were run until convergence (4 chains with a burn in of 10,000, and 10,000 sampled iterations each), with convergence being assessed using trace plots and the R hat diagnostic.(18) All numeric confounders were centered and scaled by their standard deviation, and age was adjusted for using single year of age categories.

## RESULTS

### Descriptive analysis

During the study period there were 114,820 notifications of TB in England, of which 93% (106765/114820) had their birth status recorded. Of notifications with a known birth status 27% (29096/106765) were UK born, in comparison to 33% (2634/8055) in cases with an imputed birth status. Trends in incidence rates varied by age group and UK birth status (see supplementary information). During the study period, there were 1729 UK born cases and 2797 non-UK born cases in individuals relevant to the universal schools scheme, and 1431 UK born cases and 238 non-UK born cases relevant to the targeted neonatal scheme, who fit our age criteria. Univariable evidence for differences between mean incidence rates before and after the change in BCG policy in the UK born was weak. In the non-UK born incidence rates were lower after the change in BCG policy in both the cohort relevant to the universal school-age scheme and the cohort relevant to the targeted neonatal scheme (Figure 1).

**Figure 1:**
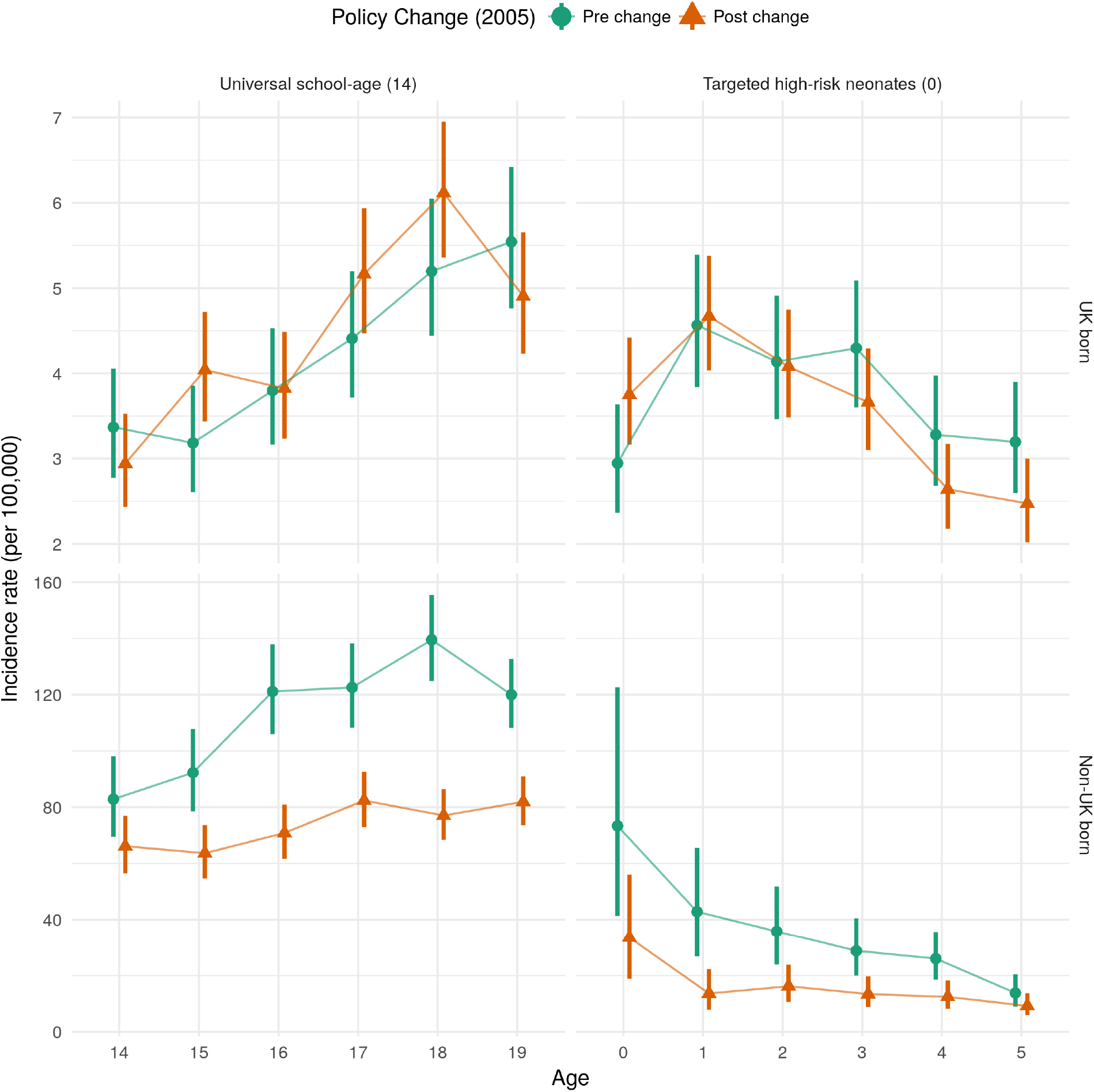
Mean incidence rates per 100,000, with 95% confidence intervals for each retrospective cohort (see table 1 for cohort definitions), stratified by the vaccination policy and UK birth status. The top and bottom panels are on different scales in order to highlight trends in incidence rates over time.

### Adjusted estimates of the effects of the change in policy on school-age children

In the UK born cohort relevant to universal vaccination there was some evidence, across all models that adjusted for age, that ending the scheme was associated with a modest increase in TB rates (Supplementary Table S2). Using the LOOIC goodness of fit criteria the best fitting model was found to be a Negative Binomial model that adjusted for the change in policy, age, and incidence rates in the UK born (Table 2). In this model there was some evidence of an assocation between the change in policy and an increase in incidence rates in those at school-age who were UK born, with an IRR of 1.08 (95%CI 0.97, 1.19). Dropping the change in policy from the model resulted in a small decrease in the LOOIC (0.52 (SE 2.63)) but the change was too small, with too large a standard error, to conclusively state that the excluding the change in policy from the model improved the quality of model fit. We found that it was important to adjust for UK born incidence rates, otherwise the impact from the change in BCG vaccination policy was over-estimated.

For the comparable non-UK born cohort who were relevant to the universal vaccination there was evidence, in the best fitting model, that ending the scheme was associated with a decrease in incidence rates (IRR: 0.74 (95%CI 0.61, 0.88)). The best fitting model was a Negative Binomial model which adjusted for the change in policy, age, incidence rates in the non-UK born, and year of eligibility as a random effect (Table 2). We found omitting change in policy from the model resulted in poorer model fit (LOOIC increase of 3.02 (SE 3.52)), suggesting that the policy change was an important factor explaining changes in incidence rates, after adjusting for other covariates. All models that adjusted for incidence rates in the UK born or non-UK born estimated similar IRRs (Supplementary Table S3).

**Table 2:** Summary table of incidence rate ratios, in the UK born and non-UK born cohorts relevant to the universal school-age scheme, using the best fitting models as determined by comparison of the LOOIC (UK born: Negative binomial model adjusting with fixed effects for the change in policy, age, and incidence rates in the UK born (Model 7 (Negative Binomial)), Non-UK born: Negative binomial model with a random intercept for year of study entry, adjusting with fixed effects for the change in policy, age, and incidence rates in the non-UK born (Model 17 (Negative Binomial))). Model terms which were not included in a given cohort are indicated using a hyphen (−).

| Variable | IRR (95% CrI)* |  |
| --- | --- | --- |
|  | UK born | Non-UK born |
| Policy change† |  |  |
| Pre-change | *Reference* | *Reference* |
| Post-change | 1.08 (0.97, 1.19) | 0.74 (0.61, 0.88) |
| Age |  |  |
| 14 | *Reference* | *Reference* |
| 15 | 1.18 (0.98, 1.42) | 1.03 (0.87, 1.22) |
| 16 | 1.24 (1.03, 1.50) | 1.25 (1.07, 1.47) |
| 17 | 1.59 (1.33, 1.91) | 1.40 (1.19, 1.63) |
| 18 | 1.92 (1.60, 2.30) | 1.47 (1.26, 1.73) |
| 19 | 1.80 (1.49, 2.17) | 1.47 (1.24, 1.73) |
| UK born incidence rate (per standard deviation) | 1.08 (1.03, 1.14) | - |
| Non-UK born incidence rate (per standard deviation) | - | 1.11 (1.03, 1.19) |
| Year of study eligibility, group level | - |  |
| Intercept (standard deviation) | - | 1.13 (1.05, 1.26) |
| Year of study eligibility, individual level | - |  |
| 2000 | - | 1.10 (0.96, 1.29) |
| 2001 | - | 1.06 (0.93, 1.24) |
| 2002 | - | 1.07 (0.94, 1.25) |
| 2003 | - | 0.90 (0.76, 1.03) |
| 2004 | - | 0.89 (0.75, 1.02) |
| 2005 | - | 0.98 (0.85, 1.12) |
| 2006 | - | 1.13 (0.99, 1.33) |
| 2007 | - | 1.04 (0.91, 1.20) |
| 2008 | - | 0.96 (0.83, 1.09) |
| 2009 | - | 0.95 (0.81, 1.08) |
| 2010 | - | 0.96 (0.82, 1.11) |
| * Incidence Rate Ratio (95% Credible Interval) |  |  |

† There was an improvement in the LOOIC score of 0.52 (SE 2.63) from dropping the change in policy from the model in the UK born cohort and a -3.02 (SE 3.52) improvement in the non-UK born cohort.

### Adjusted estimates of the effect of the change in policy in those relevant to the targeted neonatal programme

For the UK born cohort relevant to the targeted neonatal vaccination programme the evidence of an association, across all models, was mixed and credible intervals were wide compared to models for the UK born cohort relevant to the universal school-age vaccination programme (Supplementary Table S4). The best fitting model was a Poisson model which adjusted for the change in policy, age, UK born incidence rates, and year of study entry with a random effect (Table 3). In this model, there was weak evidence of an association between the change in BCG policy and an decrease in incidence rates in UK born neonates, with an IRR of 0.96 (95%CI 0.82, 1.14). There was weak evidence to suggest that dropping the change in policy from this model improved the quality of the fit, with an improvement in the LOOIC score of 0.92 (SE 1.07). This suggests that the change in policy was not an important factor for explaining incidence rates, after adjusting for covariates. Models which also adjusted for non-UK born incidence rates estimated that the change in policy was associated with no change in incidence rates in the relevant cohort of neonates.

For the comparable non-UK born cohort who were relevant to the targeted neonatal vaccination programme there was evidence, across all models, that change in policy was associated with a large decrease in incidence rates (IRR: 0.62 (95%CI 0.44, 0.88)) (Table 3 in the best fitting model). The best fitting model was a Negative Binomial model that adjusted for the change in policy, age, and non-UK born incidence rates (Table 3). All models which at least adjusted for age estimated comparable effects of the change in policy (Supplementary Table S5).

**Table 3:**
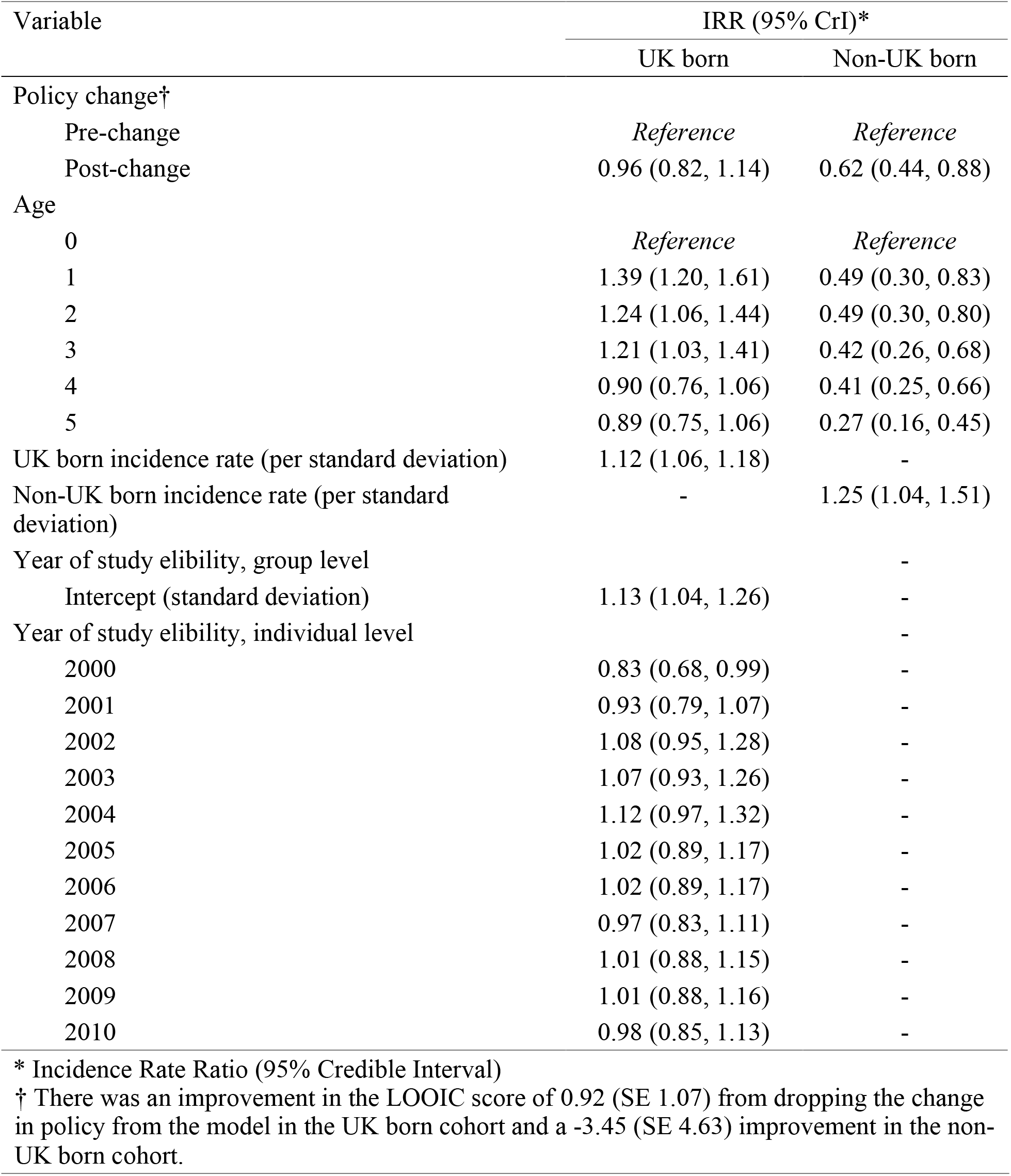
Summary table of incidence rate ratios, in the UK born and non-UK born cohorts relevant to the targeted neonatal scheme, using the best fitting models as determined by comparison of the LOOIC (UK born: Poisson model with a random intercept for year of study entry, adjusting with fixed effects for the change in policy, age, and incidence rates in the UK born (Model 16), Non-UK born: Negative binomial model adjusting with fixed effects for the change in policy, age, and incidence rates in the non-UK born (Model 8 (Negative Binomial))). Model terms which were not included in a given cohort are indicated using a hyphen (−).

| Variable | IRR (95% CrI)* |  |
| --- | --- | --- |
|  | UK born | Non-UK born |
| Policy change† |  |  |
| Pre-change | *Reference* | *Reference* |
| Post-change | 0.96 (0.82, 1.14) | 0.62 (0.44, 0.88) |
| Age |  |  |
| 0 | *Reference* | *Reference* |
| 1 | 1.39 (1.20, 1.61) | 0.49 (0.30, 0.83) |
| 2 | 1.24 (1.06, 1.44) | 0.49 (0.30, 0.80) |
| 3 | 1.21 (1.03, 1.41) | 0.42 (0.26, 0.68) |
| 4 | 0.90 (0.76, 1.06) | 0.41 (0.25, 0.66) |
| 5 | 0.89 (0.75, 1.06) | 0.27 (0.16, 0.45) |
| UK born incidence rate (per standard deviation) | 1.12 (1.06, 1.18) | - |
| Non-UK born incidence rate (per standard deviation) | - | 1.25 (1.04, 1.51) |
| Year of study eligibility, group level |  | - |
| Intercept (standard deviation) | 1.13 (1.04, 1.26) | - |
| Year of study eligibility, individual level |  | - |
| 2000 | 0.83 (0.68, 0.99) | - |
| 2001 | 0.93 (0.79, 1.07) | - |
| 2002 | 1.08 (0.95, 1.28) | - |
| 2003 | 1.07 (0.93, 1.26) | - |
| 2004 | 1.12 (0.97, 1.32) | - |
| 2005 | 1.02 (0.89, 1.17) | - |
| 2006 | 1.02 (0.89, 1.17) | - |
| 2007 | 0.97 (0.83, 1.11) | - |
| 2008 | 1.01 (0.88, 1.15) | - |
| 2009 | 1.01 (0.88, 1.16) | - |
| 2010 | 0.98 (0.85, 1.13) | - |

\* Incidence Rate Ratio (95% Credible Interval)
† There was an improvement in the LOOIC score of 0.92 (SE 1.07) from dropping the change in policy from the model in the UK born cohort and a -3.45 (SE 4.63) improvement in the non-UK born cohort.

### Magnitude of the estimated impact of the change in BCG policy

We estimate that the change in vaccination policy was associated with preventing 385 (95%CI −105, 881) cases from 2005 until the end of the study period in the directly impacted populations after 5 years of follow up (Table 4). The majority of the cases prevented were in the non-UK born, with cases increasing slightly overall in the UK born. This was due to cases increasing in the UK born at school-age, and decreasing in UK born neonates, although both these estimates had large credible intervals.

**Table 4:** Estimated number of cases prevented, from 2005 until 2015, for each vaccination programme in the study population relevant to that programme, using the best fitting model for each cohort.

| Vaccination Programme | Birth Status | Cases Prevented (95% CI*) | Notified Cases |
| --- | --- | --- | --- |
| Universal school-age (14) |  | -291 (24, -571) | 2364 |
|  | UK born | 76 (188, -26) | 969 |
|  | Non-UK born | -367 (-165, -546) | 1395 |
| Targeted high-risk neonates (0) |  | 94 (-81, 310) | 906 |
|  | UK born | 30 (-95, 173) | 800 |
|  | Non-UK born | 65 (14, 137) | 106 |
| Change in Policy† |  | 385 (-105, 881) | 3270 |
|  | UK born | -46 (-284, 199) | 1769 |
|  | Non-UK born | 431 (179, 682) | 1501 |

\*95% CI: 95% Credible Interval,
† Estimated total number of cases prevented due to the change in vaccination policy in 2005

## DISCUSSION

In the non-UK born we found evidence of an association between the change in BCG policy and a decrease in TB incidence rates in both those at school-age and neonates, after 5 years of follow up. We found some evidence that the change in BCG policy was associated with a modest increase in incidence rates in the UK born population who were relevant to the universal school-age scheme and weaker evidence of a small decrease in incidence rates in the UK born population relevant to the targeted neonatal scheme. Overall, we found that the change in policy was associated with preventing 385 (95%CI −105, 881) cases in the study population, from 2005 until the end of the study period, with the majority of the cases prevented in the non-UK born.

We were unable to estimate the impact of the change in BCG policy after 5 years post vaccination, so both our estimates of the positive and negative consequences are likely to be underestimates of the ongoing impact. Tuberculosis is a complex disease and the BCG vaccine is known to offer imperfect protection, which has been shown to vary both spatially and with time since vaccination.(19, 20) By focusing on the impact of the change in policy on the directly affected populations within a short period of time, and by employing a multi-model approach we have limited the potential impact of these issues. Our study was based on a routine observational data set (ETS), and a repeated survey (LFS) both of which may have introduced bias. Whilst the LFS is a robust data source, widely used in academic studies,(21–23) it is susceptible to sampling errors particularly in the young, and in the old, which may have biased the estimated incidence rates. As the ETS is routine surveillance system some level of missing data is inevitable. However, UK birth status is relatively complete (93% (106765/114820)) and we imputed missing values using an approach which accounted for MNAR mechanisms for the variables included in the imputation model. We were unable to adjust for known demographic risk factors for TB, notably socio-economic status,(24, 25) and ethnicity.(24–26) However, this confounding is likely to be mitigated by our use of multiple cohorts and our adjustment for incidence rates in the UK born and non-UK born. Finally, we have assumed that the effect we have estimated for the change in BCG policy is due to the changes in BCG vaccination policy as well as other associated changes in TB control policy, after adjusting for hypothesised confounders. However, there may have been additional policy changes which we have not accounted for.

Whilst little work has been done to assess the impact of the 2005 change in BCG vaccination several other studies have estimated the impact of changing BCG vaccination policy, although typically only from universal vaccination of neonates to targeted vaccination of high-risk neonates. A previous study in Sweden found that incidence rates in Swedish-born children increased after high-risk neonatal vaccination was implemented in place of a universal neonatal program, this corresponds with our finding that introducing neonatal vaccination had little impact on incidence rates in UK born neonates. Theoretical approaches have indicated that targeted vaccination of those at high-risk may be optimal in low incidence settings.(27) Our study extends this work by also considering the age of those given BCG vaccination, although we were unable to estimate the impact of a universal neonatal scheme as this has never been implemented nationally in England. It has previously been shown that targeted vaccination programmes may not reach those considered most at risk,(28) our findings may support this view as we observed only a small decrease in incidence rates in UK born neonates after the introduction of the targeted neonatal vaccination programme. Alternatively, the effectiveness of the BCG in neonates, in England, may be lower than previously thought as we only observed a small decrease in incidence rates, whilst a previous study estimated BCG coverage at 68% (95%CI 65%, 71%) amongst those eligible for the targeted neonatal vaccination programme.(29)

This study indicates that the change in England’s BCG vaccination policy was associated with a modest increase in incidence in the UK born that were relevant to the school-age vaccination programme, and with a small reduction in incidence in the UK born that were relevant to the high-risk neonatal vaccination programme, although both these estimates had wide credible intervals. We found stronger evidence of an association between the change in policy and a decrease in incidence rates in the non-UK born populations relevant to both programmes. This suggests that the change of vaccination policy to target high-risk neonates may have resulted in an increased focus on high-risk non-UK born individuals who may not have been the direct targets of the vaccination programme. Further validation is required using alternative study designs, but this result should be considered when vaccination policy changes are being considered.

It is well established that interventions against infectious diseases, such as TB, should be evaluated not only for their direct effects but also for future indirect effects via ongoing transmission. Statistical approaches such as those used in this paper are not appropriate for capturing these future indirect effects, and instead dynamic disease models should be used. In addition, this study could not evaluate the impact of the neonatal programme on the high-risk population it targets, due to a lack of reliable data. Improved coverage data for the BCG programme is required to more fully evaluate its ongoing impact.

## Acknowledgements

The authors thank the TB section at Public Health England (PHE) for maintaining the Enhanced Tuberculosis Surveillance (ETS) system; all the healthcare workers involved in data collection for the ETS.

## Contributors

SA, HC, and EBP conceived and designed the work. NJW provided guidance on the statistical methods used. SA undertook the analysis with advice from all other authors. All authors contributed to the interpretation of the data. SA wrote the first draft of the paper and all authors contributed to subsequent drafts. All authors approve the work for publication and agree to be accountable for the work.

## Accessibility of data and programming code

The code used to clean the data used in this paper can be found at: DOI:10.5281/zenodo.2551555

The code for the analysis contained in this paper can be found at: DOI:10.5281/zenodo.2583056

