## Supplementary Information for "Estimating the effect of the 2005 change in BCG policy in England: A retrospective cohort study"

### Online supplementary appendix: Estimating the effect of the 2005 change in BCG policy in England: A retrospective cohort study

Sam Abbott, Hannah Christensen, Nicky Welton, Ellen Brooks-Pollock

#### Model Definitions

*Supplementary Table S1: Full definition of each model, ordered by increasing complexity.*

| Model | Description |
|---|---|
| Model 1 | Poisson model adjusting for no fixed effects. |
| Model 2 | Poisson model adjusting with fixed effects for the change in policy. |
| Model 3 | Poisson model adjusting with fixed effects for the change in policy and incidence rates in the UK born. |
| Model 4 | Poisson model adjusting with fixed effects for the change in policy and incidence rates in the non-UK born. |
| Model 5 | Poisson model adjusting with fixed effects for the change in policy and incidence rates in the UK born and non-UK born populations. |
| Model 6 | Poisson model adjusting with fixed effects for the change in policy and age. |
| Model 7 | Poisson model adjusting with fixed effects for the change in policy, age, and incidence rates in the UK born. |
| Model 7 (Negative Binomial) | Negative binomial model adjusting with fixed effects for the change in policy, age, and incidence rates in the UK born. |
| Model 8 | Poisson model adjusting with fixed effects for the change in policy, age, and incidence rates in the non-UK born. |
| Model 8 (Negative Binomial) | Negative binomial model adjusting with fixed effects for the change in policy, age, and incidence rates in the non-UK born. |
| Model 9 | Poisson model adjusting with fixed effects for the change in policy, age, and incidence rates in the UK born and non-UK born populations. |
| Model 10 | Poisson model with a random intercept for year of study entry, adjusting for no fixed effects. |
| Model 11 | Poisson model with a random intercept for year of study entry, adjusting with fixed effects for the change in policy. |
| Model 12 | Poisson model with a random intercept for year of study entry, adjusting with fixed effects for the change in policy and incidence rates in the UK born. |
| Model 13 | Poisson model with a random intercept for year of study entry, adjusting with fixed effects for the change in policy and incidence rates in the non-UK born. |
| Model 14 | Poisson model with a random intercept for year of study entry, adjusting with fixed effects for the change in policy and incidence rates in the UK born and non-UK born populations. |
| Model 15 | Poisson model with a random intercept for year of study entry, adjusting with fixed effects for the change in policy and age. |
| Model 16 | Poisson model with a random intercept for year of study entry, adjusting with fixed effects for the change in policy, age, and incidence rates in the UK born. |
| Model 16 (Negative Binomial) | Negative binomial model with a random intercept for year of study entry, adjusting with fixed effects for the change in policy, age, and incidence rates in the UK born. |
| Model 17 | Poisson model with a random intercept for year of study entry, adjusting with fixed effects for the change in policy, age, and incidence rates in the non-UK born. |
| Model 17 (Negative Binomial) | Negative binomial model with a random intercept for year of study entry, adjusting with fixed effects for the change in policy, age, and incidence rates in the non-UK born. |
| Model 18 | Poisson model with a random intercept for year of study entry, adjusting with fixed effects for the change in policy, age, and incidence rates in the UK born and non-UK born populations. |

**Imputation of UK birth status**

As we were imputing a single variable we reformulated the imputation as a categorical prediction problem. This allowed us to use techniques from machine learning to improve the quality of our imputation, whilst also validating it using metrics supported by theory. We included year of notification, sex, age, Public Health England Centre (PHEC), occupation, ethnic group, Index of Multiple Deprivation (2010) categorised into five groups for England (IMD rank), and risk factor count (risk factors considered; drug use, homelessness, alcohol misuse/abuse and prison). However, we could not account for a possible missing not at random mechanism not captured by these covariates. To train the model we first split the data with complete UK birth status into a training set (80%), a calibration set (5%) and a test set (15%). We then fit a gradient boosted machine with the 10000 trees, early stopping (at a precision of $1e-5$, with 10 stopping rounds), a learning rate of 0.1, and a learn rate annealing of 0.99. Gradient boosted machines are a tree based method that can incorporate complex non-linear relationships and interactions. Much like a random forest model they work by ensembling a group of trees, but unlike a random forest model each tree is additive aiming to reduce the residual loss from previous trees. Once the model had been fit to the training set we performed platt scaling using the calibration data set. Our fitted imputation model had a Logloss of 0.28 on the test set, with an AUC of 0.93, both of which indicate a robust out of bag performance. We found that ethnic group was the most important variable for predicting UK birth status, followed by age and PHEC.

Using the fitted model we predicted the birth status for notifications where this was missing, using the F1 optimal threshold as our probability cut-off. It is common to impute missing values multiple times, to account for within- and between imputation variability. However, we considered this unnecessary for our analysis as the amount of missing data was small, our analysis considered only aggregate counts, our model metrics indicated a robust level of performance out of bag and any unaccounted for uncertainty would be outweighed by the uncertainty in our population denominator.(11) We found that cases with imputed birth status had a similar proportion of UK born to non-UK born cases as in the complete data (Supplementary Table S6).

*Supplementary Table S6: Comparision of UK birth status in cases with complete or imputed records.*

| Status | Birth Status | Proportion of Cases (%) | Cases |
|---|---|---|---|
| Complete | | | 106765 |
| | UK Born | 27.3 | 29096 |
| | Non-UK Born | 72.7 | 77669 |
| Imputed | | | 8055 |
| | UK Born | 32.7 | 2634 |
| | Non-UK Born | 67.3 | 5421 |

Inclusion of imputed values for UK birth status should reduce bias caused by any missing not at random mechanism captured by predictors included in the model. Graphical evaluation of UK birth status indicated that missingness has reduced over time, indicating a missing not at random mechanism. If only the complete case data then incidence rates would have reduced over the study period due to this mechanism, this may have biased our estimate of the impact of the change in policy.

**Prior choice**

Default weakly informative priors were used based on those provided by the brms package. For the population-level effects this was an improper flat prior over the reals. For both the standard deviations of group level effects and the group level intercepts this was a half student-t prior with 3 degrees of freedom and a scale parameter that depended on the standard deviation of the response after applying the link function.

**Estimating the magnitude of the estimated impact of the change in BCG policy**

We estimated the magnitude of the estimated impact from the change in BCG policy by applying the IRR estimates from the best fitting model for each cohort to the observed number of notifications from 2005 until 2015 in our study population. For the cohorts relevant to the universal school-age vaccination scheme we estimated the number of prevented cases by first aggregating cases ($C_O$) and then using the following equation,

$$C_P^i = C_O * (1 - I^i), \text{ Where } i = e,\ l,\ u.$$

Where $C_P^i$ is the predicted number of cases prevented using the mean ($e$), 2.5% bound ($l$) and 97.5% bound ($u$) of the IRR estimate $I^i$. For the cohorts relevant to the targeted high-risk neonatal scheme we used a related equation, adjusting for the fact that the populations were exposed to the scheme and we therefore had to first estimate the number of cases that would have been observed had the scheme not been implemented. After simplification this results in the following equation,

$$C_P^i = C_0 \left(\frac{1}{I^i} - 1\right), \text{ Where } i = e,\ l,\ u.$$

**Descriptive analysis of age-specific incidence rates**

From 2000 until 2012 incidence rates in the UK born remained relatively stable but have since fallen year on year. In comparison incidence rates in the non-UK born increased from 2000 until 2005, since when they have also decreased year on year. In 14-19 year old's, who were UK born, incidence rates remained relatively stable throughout the study period, except for the period between 2006 to 2009 in which they increased year on year. This trend was not observed in the non-UK born population aged 14-19, where incidence rates reached a peak in 2003, since when they have consistently declined. In those aged 0-5, who were UK born, incidence rates also increased year on year after the change in BCG policy, until 2008 since when they have declined. This does not match with the observed trend in incidence rates in the non-UK born population, aged 0-5, in which incidence rates declined steeply between 2005 and 2006, since when they have remained relatively stable (Supplementary Figure S1; Supplementary Table S7; Supplementary Table S8).

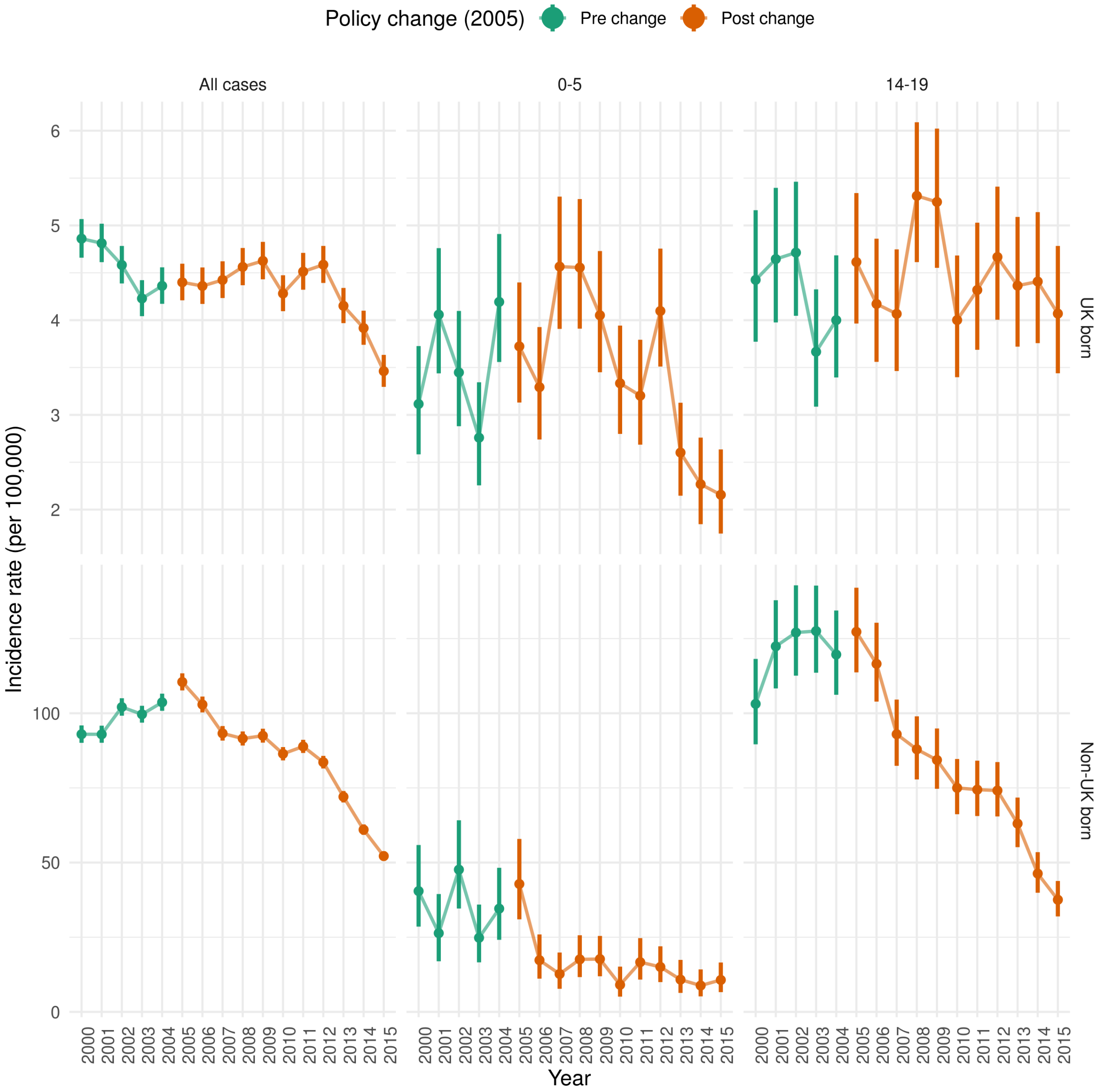


*Supplementary Figure S1: Incidence rates per 100,000 for UK born population and non-UK born population, aged 0-5 and therefore directly affected by the targeted neonatal vaccination programme, and aged 14-19 and therefore directly affected by the universal school-age scheme.*

**Incidence estimates for all cases, those aged 0-5 and those aged 14-19**

*Supplementary Table S7: Incidence rates per 100,000 in the UK born for all cases, those aged 0-5, and those aged 14-19, who were directly affected by the change in vaccination policy in 2005*

| Year eligible for vaccination | Age group | | |
|---|---|---|---|
| | All cases* | 0-5* | 14-19* |
| 2000 | 4.86 (4.66, 5.07) | 3.12 (2.58, 3.73) | 4.43 (3.77, 5.16) |
| 2001 | 4.81 (4.61, 5.02) | 4.06 (3.44, 4.76) | 4.65 (3.98, 5.40) |
| 2002 | 4.58 (4.39, 4.78) | 3.45 (2.88, 4.10) | 4.71 (4.05, 5.46) |
| 2003 | 4.23 (4.04, 4.42) | 2.76 (2.26, 3.34) | 3.67 (3.09, 4.33) |
| 2004 | 4.36 (4.17, 4.56) | 4.19 (3.56, 4.91) | 4.00 (3.40, 4.68) |
| 2005 | 4.40 (4.21, 4.60) | 3.72 (3.13, 4.40) | 4.61 (3.96, 5.34) |
| 2006 | 4.36 (4.17, 4.56) | 3.29 (2.74, 3.93) | 4.17 (3.56, 4.86) |
| 2007 | 4.42 (4.23, 4.62) | 4.57 (3.91, 5.30) | 4.07 (3.46, 4.75) |
| 2008 | 4.56 (4.37, 4.76) | 4.56 (3.91, 5.28) | 5.31 (4.61, 6.09) |
| 2009 | 4.63 (4.43, 4.83) | 4.05 (3.45, 4.73) | 5.25 (4.55, 6.02) |
| 2010 | 4.28 (4.09, 4.47) | 3.33 (2.80, 3.94) | 4.00 (3.40, 4.68) |
| 2011 | 4.51 (4.32, 4.71) | 3.20 (2.69, 3.79) | 4.32 (3.69, 5.03) |
| 2012 | 4.59 (4.39, 4.78) | 4.10 (3.51, 4.76) | 4.67 (4.00, 5.41) |
| 2013 | 4.15 (3.97, 4.34) | 2.60 (2.15, 3.13) | 4.36 (3.72, 5.09) |
| 2014 | 3.92 (3.74, 4.10) | 2.27 (1.85, 2.76) | 4.41 (3.76, 5.14) |
| 2015 | 3.46 (3.30, 3.63) | 2.16 (1.75, 2.64) | 4.07 (3.44, 4.78) |

* Incidence rate per 100,000, with 95% confidence intervals

*Supplementary Table S8: Incidence rates per 100,000 in the non-UK born for all cases, those aged 0-5, and those aged 14-19, who would have been directly affected by the change in vaccination policy in 2005 had they been UK born*

| Year eligible for vaccination | Age group | | |
|---|---|---|---|
| | All cases* | 0-5* | 14-19* |
| 2000 | 92.98 (90.10, 95.92) | 40.45 (28.56, 55.88) | 103.14 (89.60, 118.19) |
| 2001 | 92.95 (90.12, 95.84) | 26.36 (16.95, 39.47) | 122.40 (108.32, 137.85) |
| 2002 | 102.07 (99.18, 105.03) | 47.63 (34.62, 64.16) | 127.03 (112.59, 142.83) |
| 2003 | 99.65 (96.85, 102.50) | 24.81 (16.59, 35.94) | 127.53 (113.57, 142.75) |
| 2004 | 103.66 (100.82, 106.56) | 34.58 (24.13, 48.25) | 119.66 (106.18, 134.41) |
| 2005 | 110.48 (107.64, 113.37) | 42.83 (30.99, 57.91) | 127.26 (113.69, 142.04) |
| 2006 | 102.91 (100.28, 105.59) | 17.32 (11.13, 25.93) | 116.54 (103.91, 130.31) |
| 2007 | 93.26 (90.85, 95.71) | 12.69 (7.74, 19.87) | 92.99 (82.40, 104.58) |
| 2008 | 91.52 (89.19, 93.90) | 17.59 (11.66, 25.67) | 87.92 (77.84, 98.97) |
| 2009 | 92.47 (90.17, 94.82) | 17.69 (11.92, 25.44) | 84.34 (74.71, 94.90) |
| 2010 | 86.41 (84.21, 88.67) | 9.07 (5.11, 15.16) | 75.00 (66.19, 84.68) |
| 2011 | 88.88 (86.70, 91.10) | 16.65 (10.82, 24.70) | 74.41 (65.59, 84.12) |
| 2012 | 83.60 (81.51, 85.73) | 15.05 (9.97, 21.96) | 74.12 (65.45, 83.65) |
| 2013 | 72.03 (70.13, 73.97) | 10.80 (6.36, 17.41) | 63.04 (55.16, 71.77) |
| 2014 | 61.01 (59.29, 62.78) | 8.82 (5.19, 14.22) | 46.31 (39.90, 53.49) |
| 2015 | 52.18 (50.62, 53.77) | 10.69 (6.62, 16.54) | 37.55 (31.97, 43.83) |

* Incidence rate per 100,000, with 95% confidence intervals

#### **Direct effects of the change in policy on the UK born cohorts - results from all models**

*Supplementary Table S2: Comparison of models fitted to incidence rates for the UK born population that were relevant to the universal vaccination programme of those at school-age (14). Models are ordered by the goodness of fit as assessed by LOOIC, the degrees of freedom are used as a tiebreaker.*

| Model | IRR (CI 95%)* | Variable | | | | | DoF** | LPD† | LOOIC (se)†† |
|---|---|---|---|---|---|---|---|---|---|
| | | Policy Change | Age | UK born rates | Non-UK born rates | Year of study entry | | | |
| Model 7 (Negative Binomial) | 1.08 (0.97, 1.19) | Yes | Yes | Yes | No | No | 9 | -211 | 439 (10) |
| Model 7 | 1.08 (1.00, 1.17) | Yes | Yes | Yes | No | No | 8 | -211 | 443 (14) |
| Model 9 | 1.12 (1.01, 1.25) | Yes | Yes | Yes | Yes | No | 9 | -210 | 445 (14) |
| Model 16 | 1.08 (0.97, 1.21) | Yes | Yes | Yes | No | Yes | 20 | -207 | 445 (14) |
| Model 18 | 1.12 (0.97, 1.28) | Yes | Yes | Yes | Yes | Yes | 21 | -207 | 447 (15) |
| Model 8 | 1.16 (1.04, 1.29) | Yes | Yes | No | Yes | No | 8 | -213 | 449 (17) |
| Model 6 | 1.06 (0.98, 1.15) | Yes | Yes | No | No | No | 7 | -215 | 452 (17) |
| Model 17 | 1.15 (1.00, 1.32) | Yes | Yes | No | Yes | Yes | 20 | -209 | 452 (17) |
| Model 15 | 1.06 (0.94, 1.20) | Yes | Yes | No | No | Yes | 19 | -209 | 453 (17) |
| Model 1 | 0.00 (0.00, 0.00) | No | No | No | No | No | 1 | -254 | 513 (26) |
| Model 2 | 1.06 (0.98, 1.14) | Yes | No | No | No | No | 2 | -252 | 515 (25) |
| Model 4 | 1.00 (0.90, 1.10) | Yes | No | No | Yes | No | 3 | -251 | 516 (25) |
| Model 3 | 1.06 (0.98, 1.15) | Yes | No | Yes | No | No | 3 | -252 | 518 (26) |
| Model 5 | 0.98 (0.89, 1.09) | Yes | No | Yes | Yes | No | 4 | -249 | 518 (24) |
| Model 13 | 0.94 (0.78, 1.12) | Yes | No | No | Yes | Yes | 15 | -237 | 518 (27) |
| Model 10 | 0.00 (0.00, 0.00) | No | No | No | No | Yes | 13 | -244 | 521 (28) |
| Model 11 | 1.06 (0.94, 1.20) | Yes | No | No | No | Yes | 14 | -244 | 522 (28) |
| Model 14 | 0.93 (0.78, 1.11) | Yes | No | Yes | Yes | Yes | 16 | -236 | 522 (27) |
| Model 12 | 1.06 (0.93, 1.20) | Yes | No | Yes | No | Yes | 15 | -243 | 526 (28) |

* Incidence Rate Ratio, with 95% credible intervals,
** Degrees of Freedom,
† Computed log pointwise predictive density,
†† Leave one out information criterion, with standard error

*Supplementary Table S4: Comparison of models fitted to incidence rates for the UK born population that were elgible to the targeted vaccination programme of neonates. Models are ordered by the goodness of fit as assessed by LOOIC, the degrees of freedom are used as a tiebreaker.*

| Model | IRR (CI 95%)* | Variable | | | | | DoF** | LPD† | LOOIC (se)†† |
|---|---|---|---|---|---|---|---|---|---|
| | | Policy Change | Age | UK born rates | Non-UK born rates | Year of study entry | | | |
| Model 16 | 0.96 (0.82, 1.14) | Yes | Yes | Yes | No | Yes | 20 | -192 | 415 (12) |
| Model 16 (Negative Binomial) | 0.96 (0.82, 1.13) | Yes | Yes | Yes | No | Yes | 21 | -196 | 415 (10) |
| Model 16 (Negative Binomial) | 0.96 (0.82, 1.13) | Yes | Yes | Yes | No | Yes | 21 | -196 | 415 (10) |
| Model 18 | 0.99 (0.82, 1.18) | Yes | Yes | Yes | Yes | Yes | 21 | -192 | 417 (13) |
| Model 7 | 0.96 (0.88, 1.05) | Yes | Yes | Yes | No | No | 8 | -200 | 420 (15) |
| Model 9 | 1.00 (0.89, 1.12) | Yes | Yes | Yes | Yes | No | 9 | -200 | 422 (15) |
| Model 8 | 1.02 (0.91, 1.15) | Yes | Yes | No | Yes | No | 8 | -203 | 427 (16) |
| Model 6 | 0.95 (0.87, 1.03) | Yes | Yes | No | No | No | 7 | -204 | 428 (16) |
| Model 15 | 0.95 (0.83, 1.09) | Yes | Yes | No | No | Yes | 19 | -198 | 428 (14) |
| Model 17 | 1.02 (0.87, 1.20) | Yes | Yes | No | Yes | Yes | 20 | -198 | 429 (14) |
| Model 14 | 1.10 (0.92, 1.33) | Yes | No | Yes | Yes | Yes | 16 | -206 | 442 (16) |
| Model 5 | 1.08 (0.97, 1.21) | Yes | No | Yes | Yes | No | 4 | -216 | 445 (18) |
| Model 12 | 0.98 (0.83, 1.15) | Yes | No | Yes | No | Yes | 15 | -209 | 448 (17) |
| Model 4 | 1.12 (1.00, 1.24) | Yes | No | No | Yes | No | 3 | -219 | 449 (18) |
| Model 3 | 0.97 (0.89, 1.06) | Yes | No | Yes | No | No | 3 | -219 | 450 (19) |
| Model 13 | 1.14 (0.97, 1.35) | Yes | No | No | Yes | Yes | 15 | -211 | 452 (16) |
| Model 1 | 0.00 (0.00, 0.00) | No | No | No | No | No | 1 | -229 | 462 (21) |
| Model 2 | 0.95 (0.87, 1.03) | Yes | No | No | No | No | 2 | -228 | 463 (20) |
| Model 10 | 0.00 (0.00, 0.00) | No | No | No | No | Yes | 13 | -220 | 466 (19) |
| Model 11 | 0.95 (0.83, 1.09) | Yes | No | No | No | Yes | 14 | -219 | 467 (19) |

* Incidence Rate Ratio, with 95% credible intervals,
** Degrees of Freedom,
† Computed log pointwise predictive density,
†† Leave one out information criterion, with standard error

#### **Direct effects of the change in policy on the non-UK born cohorts - results from all models**

*Supplementary Table S3: Comparison of models fitted to incidence rates for the non-UK born population that were eligible to the universal vaccination programme of those at school-age (14). Models are ordered by the goodness of fit as assessed by LOOIC, the degrees of freedom are used as a tiebreaker.*

| Model | IRR (CI 95%)* | Variable | | | | | DoF** | LPD† | LOOIC (se)†† |
|---|---|---|---|---|---|---|---|---|---|
| | | Policy Change | Age | UK born rates | Non-UK born rates | Year of study entry | | | |
| Model 17 (Negative Binomial) | 0.74 (0.61, 0.88) | Yes | Yes | No | Yes | Yes | 21 | -228 | 483 (10) |
| Model 17 | 0.74 (0.62, 0.87) | Yes | Yes | No | Yes | Yes | 20 | -223 | 492 (16) |
| Model 18 | 0.73 (0.61, 0.87) | Yes | Yes | Yes | Yes | Yes | 21 | -222 | 493 (16) |
| Model 15 | 0.64 (0.53, 0.78) | Yes | Yes | No | No | Yes | 19 | -224 | 496 (18) |
| Model 16 | 0.65 (0.54, 0.78) | Yes | Yes | Yes | No | Yes | 20 | -223 | 496 (17) |
| Model 8 | 0.79 (0.73, 0.86) | Yes | Yes | No | Yes | No | 8 | -239 | 507 (20) |
| Model 9 | 0.79 (0.72, 0.86) | Yes | Yes | Yes | Yes | No | 9 | -238 | 511 (20) |
| Model 11 | 0.64 (0.52, 0.78) | Yes | No | No | No | Yes | 14 | -241 | 522 (22) |
| Model 10 | 0.00 (0.00, 0.00) | No | No | No | No | Yes | 13 | -241 | 523 (22) |
| Model 12 | 0.64 (0.53, 0.79) | Yes | No | Yes | No | Yes | 15 | -241 | 525 (22) |
| Model 13 | 0.64 (0.52, 0.79) | Yes | No | No | Yes | Yes | 15 | -241 | 526 (23) |
| Model 14 | 0.64 (0.52, 0.79) | Yes | No | Yes | Yes | Yes | 16 | -241 | 530 (23) |
| Model 7 | 0.66 (0.62, 0.70) | Yes | Yes | Yes | No | No | 8 | -248 | 532 (23) |
| Model 6 | 0.65 (0.61, 0.69) | Yes | Yes | No | No | No | 7 | -253 | 539 (27) |
| Model 4 | 0.70 (0.65, 0.76) | Yes | No | No | Yes | No | 3 | -270 | 556 (31) |
| Model 5 | 0.70 (0.64, 0.76) | Yes | No | Yes | Yes | No | 4 | -270 | 559 (31) |
| Model 2 | 0.65 (0.61, 0.69) | Yes | No | No | No | No | 2 | -275 | 561 (33) |
| Model 3 | 0.65 (0.61, 0.69) | Yes | No | Yes | No | No | 3 | -273 | 561 (32) |
| Model 1 | 0.00 (0.00, 0.00) | No | No | No | No | No | 1 | -341 | 692 (51) |

* Incidence Rate Ratio, with 95% credible intervals,
** Degrees of Freedom,
† Computed log pointwise predictive density,
†† Leave one out information criterion, with standard error

*Supplementary Table S5: Comparison of models fitted to incidence rates for the non-UK born population that were revelant to the targeted vaccination programme of neonates. Models are ordered by the goodness of fit as assessed by LOOIC, the degrees of freedom are used as a tiebreaker.*

| Model | IRR (CI 95%)* | Variable | | | | | DoF** | LPD† | LOOIC (se)†† |
|---|---|---|---|---|---|---|---|---|---|
| | | Policy Change | Age | UK born rates | Non-UK born rates | Year of study entry | | | |
| Model 8 (Negative Binomial) | 0.62 (0.44, 0.88) | Yes | Yes | No | Yes | No | 9 | -138 | 293 (15) |
| Model 8 | 0.64 (0.47, 0.86) | Yes | Yes | No | Yes | No | 8 | -137 | 295 (18) |
| Model 9 | 0.62 (0.45, 0.85) | Yes | Yes | Yes | Yes | No | 9 | -137 | 297 (18) |
| Model 6 | 0.47 (0.38, 0.58) | Yes | Yes | No | No | No | 7 | -139 | 298 (19) |
| Model 7 | 0.48 (0.39, 0.60) | Yes | Yes | Yes | No | No | 8 | -139 | 298 (19) |
| Model 17 | 0.63 (0.44, 0.89) | Yes | Yes | No | Yes | Yes | 20 | -135 | 298 (18) |
| Model 18 | 0.61 (0.42, 0.87) | Yes | Yes | Yes | Yes | Yes | 21 | -135 | 300 (18) |
| Model 15 | 0.47 (0.35, 0.62) | Yes | Yes | No | No | Yes | 19 | -136 | 301 (20) |
| Model 16 | 0.48 (0.36, 0.63) | Yes | Yes | Yes | No | Yes | 20 | -136 | 301 (19) |
| Model 4 | 0.82 (0.61, 1.10) | Yes | No | No | Yes | No | 3 | -147 | 304 (17) |
| Model 5 | 0.78 (0.58, 1.06) | Yes | No | Yes | Yes | No | 4 | -147 | 306 (18) |
| Model 13 | 0.83 (0.59, 1.16) | Yes | No | No | Yes | Yes | 15 | -145 | 308 (18) |
| Model 14 | 0.78 (0.55, 1.12) | Yes | No | Yes | Yes | Yes | 16 | -144 | 310 (19) |
| Model 3 | 0.52 (0.42, 0.64) | Yes | No | Yes | No | No | 3 | -152 | 314 (22) |
| Model 12 | 0.51 (0.38, 0.69) | Yes | No | Yes | No | Yes | 15 | -148 | 317 (23) |
| Model 2 | 0.49 (0.40, 0.61) | Yes | No | No | No | No | 2 | -156 | 319 (22) |
| Model 11 | 0.49 (0.37, 0.65) | Yes | No | No | No | Yes | 14 | -152 | 322 (23) |
| Model 10 | 0.00 (0.00, 0.00) | No | No | No | No | Yes | 13 | -150 | 330 (25) |
| Model 1 | 0.00 (0.00, 0.00) | No | No | No | No | No | 1 | -171 | 346 (27) |

* Incidence Rate Ratio, with 95% credible intervals,
** Degrees of Freedom,
† Computed log pointwise predictive density,
†† Leave one out information criterion, with standard error
